# Identifying gaps between scientific and local knowledge in climate change adaptation for northern European agriculture

**DOI:** 10.1101/2025.06.23.661058

**Authors:** Kristina Blennow, Georg Carlsson, Laura Grenville-Briggs, Per Hansson, Åsa Lankinen

**Affiliations:** Department of Landscape Architecture, Planning and Management, Swedish University of Agricultural Sciences; Department of Physical Geography and Ecosystem Science, Lund University; Department of Biosystems and Technology, Swedish University of Agricultural Sciences; Department of Plant Protection Biology, Swedish University of Agricultural Sciences; Swedish Centre for Agricultural Business Management, Department of People and Society, Swedish University of Agricultural Sciences

**Keywords:** 2*018 drought*, *temperate agriculture*, *climate change adaptation*, *crisis management*, *crop production*, *local knowledge*

## Abstract

Climate change impacts agriculture in complex and regionally specific ways. In temperate regions such as southern Sweden, it presents both opportunities, such as longer growing seasons, and challenges, including increased drought and heat stress. This study examined how Swedish agriculture is adapting to climate change, using the extreme 2018 drought and heatwave as a critical case to explore both immediate responses and longer-term preparedness in crop production systems. We compared scientific and local knowledge through a systematic scoping review of peer-reviewed literature from northern Europe, a workshop with project scientists, and a participatory workshop with farmers. The study aimed to identify gaps between scientific and local understandings of climate change adaptation, particularly in how extreme events influence decision-making and preparedness. Using a transdisciplinary, problem-feeding approach, we compared how farmers and scientists conceptualise crisis response and long-term adaptation, employing influence diagrams to visualise their understandings. Our review found that much scientific work focuses on short-term coping strategies, such as feed substitution or altered irrigation, while longer-term adaptation remains underexplored. In contrast, farmers emphasized context specific actions grounded in their farm operations, highlighting socio-economic and administrative constraints, whereas scientists tended to propose generalisable technology-oriented strategies. These differences underscore the importance of integrating both experiential and scientific knowledge to co-develop more effective adaptation responses. The study lays the foundation for a targeted, large-scale survey to explore farmer motivations, perceptions of risk and opportunity, and adaptive capacities across diverse contexts. Our findings highlight the need to move beyond reactive measures and investigate anticipatory strategies that foster and enable adaptive capacity. Here we demonstrate that supporting long-term adaptation in agriculture will require inclusive, evidence-based approaches that incorporate perspectives from farmers, advisors, scientists, and public authorities alike.

## 1. Introduction

Agriculture is critical for food security, rural livelihoods, and economic development. However, climate change presents both opportunities such as extended growing seasons, and challenges, including droughts and heatwaves. Adaptation is essential to mitigate losses and leverage emerging opportunities, making it crucial to understand farmers’ needs and motivations to support adaptation.

In northern Europe the climate is projected to shift towards longer growing seasons, milder winters, increased precipitation, and more frequent heatwaves and droughts (Sjökvist et al. 2025). Although these changes may enable higher crop yields and greater crop diversity (Wiréhn 2018), they also increase the risk of soil erosion, pest outbreaks, and harvest disruptions (de Toro et al. 2015; Fischer et al. 2021).

An example of these climate-induced challenges is the 2018 heatwaves and drought across northern Europe, which had severe consequences for agriculture (Sinclair et al. 2019; Toreti et al. 2019) (Figure 1). Long-term temperature records, including a 263-year dataset from Stockholm and a comprehensive 150-year dataset for Sweden as a whole, highlighted an exceptionally high mean monthly temperature in May as a defining feature of this event (Wilcke et al., 2020). Understanding the diverse implications of such phenomena is crucial for guiding adaptive measures to address future climate change.

**Figure 1.**
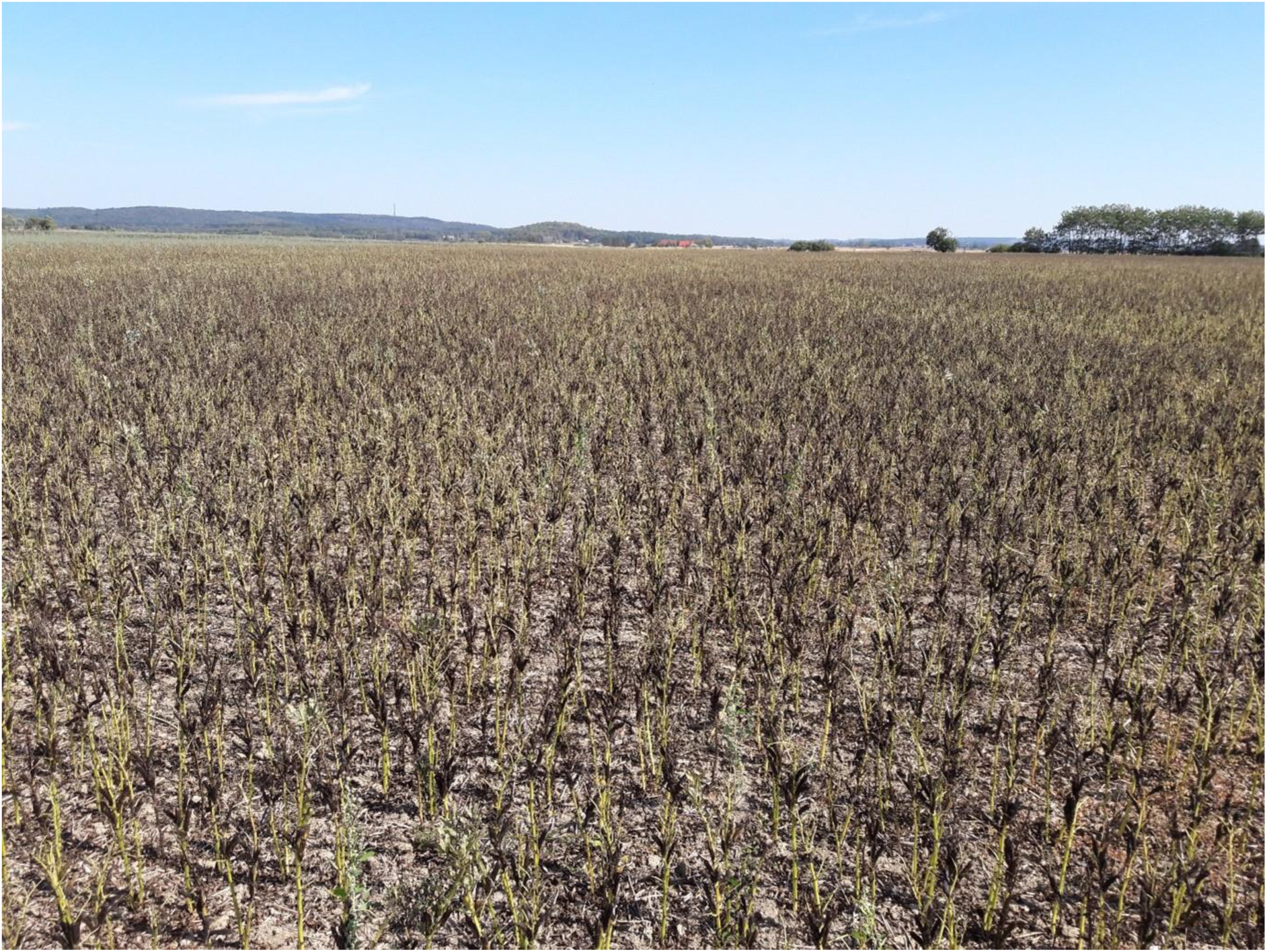
Visible signs of drought and heat stress in a faba bean (*Vicia faba* L.) field during the 2018 drought and heatwave in Sweden (31 July 2018). The photo illustrates common physiological responses such as low plant height, wilting and early maturation, highlighting the impact of extreme weather on crop performance. Photo: Georg Carlsson

This convergence of extreme weather events had profound consequences, particularly for agriculture across northern Europe (JRC, 2018). The simultaneous occurrence of droughts and heatwaves exacerbated their individual impacts, creating significant challenges to agricultural productivity, water resources, and ecosystems (Arreyndip, 2021; Buras et al. 2020; Grusson et al. 2021). For instance, cereal yields in Sweden dropped by 43%, compared to the five-year average, while reductions in fodder forced many farmers to use harvested winter fodder for supplementary feeding during the summer (Statistics Sweden, 2019).

While farmers possess local expertise, gaps in scientific understanding can hinder the adoption of more complex practices (Ingram, 2008). For instance, farmers may be familiar with their local soil conditions but lack knowledge of advanced soil management techniques (Ingram, 2008). A recent study on Öland, Sweden, found that limited understanding of the carrying capacity of land and water resources impedes the adoption of climate-smart farming techniques (Ibrahim & Johansson, 2021).

Pest management also faces challenges from climate change, with rising temperatures enabling spread of pests from southern Europe (Bebber et al 2013; Gregory et al. 2009). Current integrated pest management (IPM) strategies, which are central to the EU’s directive on sustainable use of pesticides, lack adaptation tools for climate-induced changes (Heeb et al. 2019). Developing climate-smart pest management (CSPM) is vital, yet adoption by farmers remains limited (Gvozdenac et al 2022). Effectively addressing these challenges requires bridging the gap between scientific research and farmers’ local knowledge.

Historically, agricultural knowledge transfer followed a top-down model, i.e. considering knowledge from authorities and scientists, while marginalising farmers’ insights (see Ingram 2014). However, contemporary approaches recognise that sustainable practices often stem from local knowledge, necessitating participatory, two-way communication between researchers and farmers (Fischhoff 1995, Hamilton-Webb et al., 2017; Labeyrie et al., 2021). Studies have shown strong belief in the local impacts of climate change to be a precondition for adaptation measures among land managers (Blennow et al. 2012; Blennow et al. 2020). Additionally, adaptation requires a detailed understanding of local impacts of climate change and the ability for decision-making agents to apply this knowledge in their specific contexts (Semenza et al., 2011; Blennow & Persson, 2021). Recent findings indicate that both negative and positive experiences of climate change can promote adaptation decisions (Blennow and Persson 2021; Blennow et al. 2021).

Local “proven experience” can be as influential as scientific evidence in shaping people’s worldviews, through experiential knowledge (Persson et al. 2019; Persson et al. 2022). Comparing local “proven experience” with expert knowledge highlights the gap between “what decision-makers know” and “what they need to know”. Understanding this gap is crucial for improving communication effectiveness (Fischhoff 2013; Palmér et al. 2023).

### 1.1 Aims

This study aims to compile and analyse scientific and local knowledge about the effects of the 2018 drought-heatwave and about possible ways for agriculture to adapt to a changing climate. More specifically, the overall aim is to identify gaps between scientific and local knowledge about adaptation, not only to drought and heat but also to climate change more broadly, with a particular focus on crop production. The study also seeks to identify knowledge gaps within the scientific literature itself. The study uses knowledge from a systematic scoping review of scientific literature, knowledge of scientists in the project, and contributions from a workshop with farmers. The focus is on capturing firsthand experiences and local insights regarding the 2018 drought-heat event and extend this to opportunities and challenges for climate change adaptation in temperate agriculture with Sweden as an example. Employing a problem-feeding interdisciplinary and transdisciplinary approach (Thorén and Persson 2013; Persson et al. 2019), the study identifies knowledge gaps and lays the foundation for developing guidelines for effective communications that help integrate scientific evidence with local insights. The results are expected to improve the understanding of how future extreme events may affect agricultural systems and how farmers respond to such challenges.

## 2. Materials and Methods

### 2.1 Systematic scoping review

The RepOrting standings for Systematic Evidence Synthesis: pro-forma review protocol (ROSES) was used as the guideline for the systematic scoping review. This gold standard method supports the production of high-quality Systematic Scoping reviews and is particularly applicable to topics outside medical research fields (Haddaway et al 2018). Using the ROSES process, we first formulated our research question as “What are the potential impacts of climate change on Swedish agriculture and how can knowledge on these impacts guide adaptation?”.

Later we narrowed down to focus on the compound drought and heat event in 2018 and its impacts on agriculture and options for management that support agriculture’s adaptation to climate change. Based on our research question, we then proceeded to the literature search, which was comprised of three main processes: identification, screening and eligibility.

#### 2.1.1 Identification

In the first step, we identified five main areas for the search process (related to climate change adaptation, in: agriculture, horticulture or farming, related to Sweden, or Swedish conditions, and empirical or modelling studies. Next these keyword areas were enriched to allow us to retrieve as many relevant articles for the systematic scoping review as possible. Synonyms, related terms and variations on the original keywords were sought, and the final search string was selected for the screening process (Table 1).

**Table 1.** The search string.

| Table 2. The search string. |  |
| --- | --- |
| #<br>1 | TS=((climate NEAR/3 (change* OR cris* OR breakdown OR emergenc*)) OR "global warming" OR "global heating" OR (extreme NEAR/2 (climat* OR weather*))) |
| #<br>2 | TS=((agricultur* OR farm* OR "crop production" OR cultivation OR horticulture OR agronom* OR husbandry OR tillage OR fertilization* OR fertilisation* OR irrigation* OR drainage OR (choice NEAR/3 (crop* OR variety)) OR "spring crop*" OR "winter crop*" OR "pest management" OR IPM OR "disease control*" OR "invasive specie*") AND (adapt* OR change* OR accomodat* OR adjust* OR alter* OR modifi* OR reshap* OR revis* OR conver* OR mitigat* OR manag*)) |
| #<br>3 | TS=(Swed* OR Norw* OR Finland OR finnish OR Denmark OR danish OR Iceland* OR Ireland OR irish OR UK OR united kingdom OR england OR scotland OR scottish OR wales OR welsh OR great britain OR british OR Netherlands OR Holland OR dutch OR Belgi* OR German* OR Poland OR Lithuania* OR Latvia* OR Estonia* OR Belarus* OR Russia* OR Canad* OR Montana OR Indiana OR Illinois OR Wyoming OR Minnesota OR Ohio OR Colorado OR Maine OR "New Hampshire" OR Virginia OR Iowa OR Massachusetts OR Pennsylvania OR "Washington state" OR Oregon OR "North Dakota" OR "South Dakota" OR "New Jersey" OR Vermont OR "West Virginia" OR "North Carolina" OR "South Carolina" OR Missouri OR Idaho OR Tennessee OR Kentucky OR Maryland OR Delaware) |
| #<br>4 | (#1 AND #2 AND #3 AND PY=(2023 OR 2022 OR 2021 OR 2020 OR 2019 OR 2018) AND (LA=("ENGLISH") AND DT=("ARTICLE")) |

The search process involved two primary databases, Scopus and Web of Science (core collection), which was supplemented by searches in CABI databases, and a literature search was conducted in November 2023. From this process, 19,317 records were identified, which resulted in 11,653 records after duplicates were removed that were used for the screening process (Figure 2).

**Figure 2.**
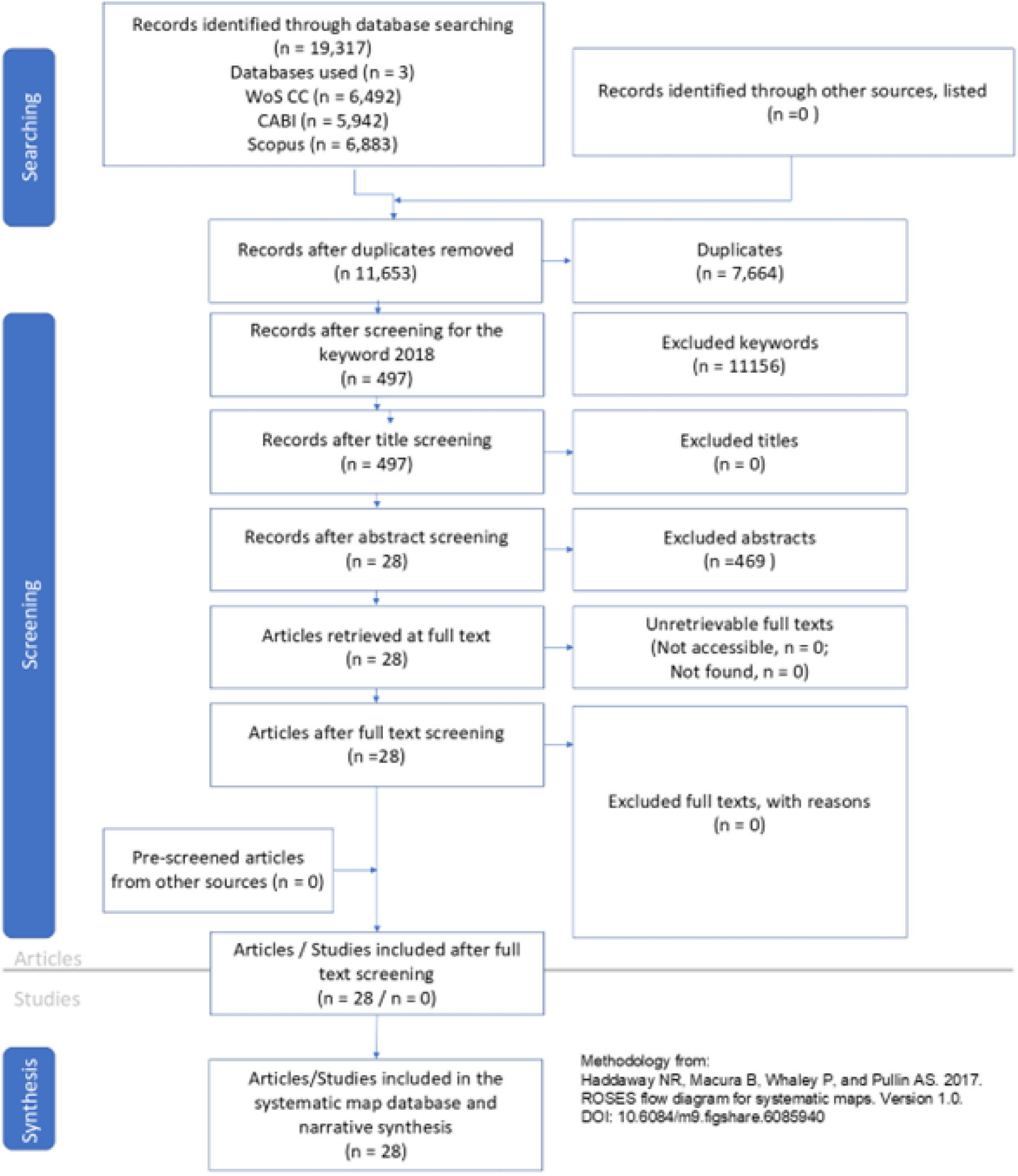
ROSES flowchart detailing the searching, screening and synthesis methodology used in this study, with the number of records used, or excluded in each step shown in the blue boxes (methodology from Haddaway et al 2017).

#### 2.1.2 Screening

In the second process, a set of inclusion and exclusion criteria were formulated for screening of the articles to ensure relevance to our research question. Firstly, it was essential to ensure that the context of the studies related to the extreme weather events during 2018. Secondly, we determined that the focus should be on studies in agriculture, horticulture or farming in general and related to adaptation. This resulted in the following inclusion and exclusion criteria.

Our inclusion criteria were:

- related to the 2018 European drought and heat extreme events related to farming, agriculture, horticulture,
- related to Swedish conditions (but study not necessarily conducted in Sweden)
- biogeophysics OR management
- empirical AND/OR modelling

Our exclusion criteria were:

- only mitigation
- only outside of the geographical areas in search strings
- only on farmer’s perceptions
- only economics
- only policies
- only information
- only communication

#### 2.1.3 Eligibility

For the third step, the relevancy of each article was manually screened by reading the title and abstract, and if required the content. This eligibility step was performed in the online systematic review tool Rayyan (Ouzzani et al 2016). After screening for the keyword 2018, 497 articles remained. Each of these 497 articles was screened at the title and abstract level by four researchers independently. The results were then compared and consolidated, leaving 28 articles retrieved at full text for inclusion within the scoping review. These articles were then reviewed in full and the findings in 17 of these are presented and discussed in detail.

### 2.2 Identification of knowledge gaps between farmers and scientists

To explore potential knowledge gaps between scientists and farmers, we employed a problem-feeding approach, whereby real-world problems are fed to science (Thorén and Persson 2013; Persson et al. 2019). Project scientists were asked to depict their understanding of relevant crisis management strategies for the 2018 heat and drought event, as well as their views on measures for adapting crop cultivation to climate change. These understandings were expressed through influence diagrams (Wood et al., 2012), which illustrate believed causal relationships and key factors influencing decision-making. The influence diagrams were constructed using Analytica by Lumina Decision Systems (2025).

To identify knowledge gaps, we compared the influence diagrams constructed separately by scientists and farmers. Discrepancies between the diagrams were interpreted as indicative of divergent mental maps or areas where communication and mutual understanding may be lacking.

#### 2.2.1 Influence Diagram Methodology

Influence diagrams were employed to explore the factors shaping climate change adaptationdecisions for crop production among farmers in Scania (Wood et al. 2012). Two types of influence diagrams were developed:

1. Scientific Perspective: Two researcher-led (the authors of this paper) influence diagrams were constructed based on expert knowledge and insights from the scoping review, which focused on the 2018 heatwave and drought event. One influence diagram focused on crisis management during a heat and drought event, which would not necessarily be motivated by a changing climate but rather would be dealing with an ongoing crisis. In the second influence diagram, the scientists contributed their expertise regarding heat and drought and other climate change-related impacts and potential adaptation measures. This diagram captured the primary climate change-related impacts and potential adaptation measures from the scientists’ perspectives.
2. Farmer Perspective: The second type of influence diagram was developed based on a workshop together with farmers conducted on 4 November 2024 at the Swedish University of Agricultural Sciences (SLU) in Alnarp. During the workshop, participating farmers identified crisis management during the 2018 heat and drought event as well as key climate change-related impacts, adaptation measures, and constraints affecting climate change adaptation in crop production. The discussion began with the 2018 drought and heatwave event and expanded to cover all aspects of climate change in relation to crop production. Consequently, the influence diagram was split into two: one diagram depicted crisis management during the 2018 heatwave and drought event, and the other diagram represented the farmers’ perspectives on climate change-related impacts, adaptation measures, and constraints for implementation of adaptation to climate change. The workshop followed a participatory approach, ensuring that diverse perspectives were captured.

#### 2.2.2 Farmers participating in the workshop

Ten farmers were invited to participate, selected from a farmer and advisor network established at the Swedish Centre for Agricultural Business Management at SLU. All participating farmers are active in Scania County, situated in southern Sweden (Figure 2). The region is characterised by its extensive agricultural land, which covers 45% of the total area, and a high concentration of agricultural enterprises (Statistics Sweden, 2025). Scania has the most fertile arable land in Sweden, with agricultural production primarily focused on annual crops, particularly cereals (Swedish Board of Agriculture, 2025). In 2024, Scania’s registered arable land totalled 433,000 ha, accounting for 17% of Sweden’s total arable land area of 2.5 million ha, excluding pastureland (Swedish Board of Agriculture, 2025).

#### 2.2.3 Comparative Analysis of Influence Diagrams

The two types of influence diagrams on crisis management and climate change adaptation, respectively, were compared to identify gaps between scientistś knowledge and farmers’ knowledge and practical experiences in crisis management, as well as adaptation and barriers to the adoption of climate change adaptation measures. Key areas of divergence were analysed to highlight:

- differences in crisis management,
- differences in perceived risk, opportunities, and adaptation priorities, and
- constraints on adaptation at the farm level (e.g., economic, regulatory, knowledge-based).

This comparative analysis provides insights into how scientific knowledge align with, or differ from, farmers’ real-world decision-making, thereby feeding real world problems to science and informing strategies to improve adaptation support for Swedish agriculture.

## 3. Results

### 3.1. Systematic scoping review

#### 3.1.1 Impacts of drought and heat on agriculture

We found 17 relevant articles that studied the effects of drought and heat on agricultural crops, at least partly involving 2018. The studies were conducted in Germany, the Netherlands, Lithuania, Poland, the UK and Sweden. The crops investigated included winter and spring wheat, rye and winter cereal, barley, oilseed rape, potato, sugar beet, onion, celeriac, faba bean, silage maize, grass and forage crops and hemp. Several of these studies also investigated potential adaptation measures (see 3.1.2).

A study from the UK that investigated the overall impact of the heat and drought in 2018 showed that most impacts were negative. In particular growth and development of crops was poor and lead to reduced yields for both food and forage crops (Holman et al. 2021). Some positive impacts included increased prices, less pests and diseases and improved soil conditions for farm operations. However, these positive factors did not outweigh the reduced yields and increased costs related to the drought.

Long-term weather and yield data (19 - 60 years) that also included 2018, confirmed that yield was negatively affected by the drought in 2018 in wheat, rye, barley, oilseed rape, potato, sugar beet, onion, silage maize and grasslands (Emadodin et al. 2021; Reinermann et al. 2019; van Oort et al. 2023; Webber et al. 2020). Webber et al. (2020) suggested that the yield loss in 2018 in barley, oilseed rape and silage maize was mainly caused by drought stress, while wheat also appeared to suffer from heat stress (see also Nehe et al. 2023). In van Oort et al. (2023), it was found that crops with deeper root systems tolerate drought better. Onion and grasses were particularly negatively affected by the 2018 drought. Moreover, in this study a comparison between irrigated and non-irrigated fields were conducted, which showed no yield loss when a good irrigation system was available. A study on the effect of experimentally reduced rainfall by 50% in semi-natural grasslands in the UK over three years (2016-2018), suggested that these types of permanent pastures are less affected by drought but still a drought in spring 2017 reduced biomass production (Ayling et al. 2021). The long-term patterns investigated in Reinermann et al. (2019) also indicated that long-lasting winters with late phenological onsets and frosts may have an important impact on yield in combination with drought.

Other studies that investigated effects of the drought in 2018 focused on effects of soil nitrogen (Klages et al. 2020; Statkevičiūtė et al. 2022), cultivar differences (Nehe et al. 2023; Statkevičiūtė et al. 2022), and pests and beneficial organisms (Nehe et al. 2023; Raderschall et al. 2022). Klages et al. (2020) showed that there were higher surpluses of nitrogen in the soil due to the drought and reduced biomass production, which may affect leakage from the field into the ground water. The spring after the drought (2019) there was still elevated concentrations of nitrogen in the soil, reducing the need for fertilisation. Studies on cultivar differences were conducted in wheat. In a study in Sweden 2016-2020 environmental factors, in particular heat rather than drought, showed stronger effects than cultivar on yield in spring wheat, but heat-tolerant cultivars could be identified (Nehe et al. 2023). In winter wheat in Lithuania 2018-2019, genotype had the strongest influence on gluten protein characteristics independent of nitrogen fertilisation or temperature (Statkevičiūtė et al. 2022). Likewise, cultivars differed in bread-baking quality. Only two of our studies considered pests or beneficial organisms. In the Swedish study on spring wheat cultivars, yield loss caused by fungal diseases was less than 1% in 2018 compared to 13% the year of the study with the highest loss (Nehe et al. 2023). In another study in Sweden on faba bean with or without annual flower strips in 2018, it was seen that the flower strips did not increase wild pollinator densities but attracted more natural enemies (ground-dwelling predators) (Raderschall et al. 2022). Thus, despite the drought flower strips may still provide beneficial habitats to natural enemies.

#### 3.1.2 Adaptation (Agricultural Planning and Management)

The reviewed studies included analyses of both short-term coping strategies, *i.e.* handling the crisis caused by drastic reduction in crop production or acute water shortage during the 2018 season, and longer-term adaptation measures. A study from the UK (Holman et al. 2021) found that short-term coping strategies were the most common responses, while measures for long-term adaptation or to improve adaptive capacity were almost lacking. As short-term coping strategies, modifications of planting and harvesting practises (*e.g.* changing the time of day for planting or harvesting potato) and maximizing the irrigation capacity were common responses (Holman et al. 2021). Changing the management of crop residues were also mentioned, it was common to harvest straw for use as animal feed, bedding or selling to other farmers, instead of incorporating straw to the soil (which would be the standard practice in a year with average rainfall and temperature) (Holman et al. 2021). Among livestock farmers, reducing the animal herd size (selling or slaughtering animals), changing the management of pastures (*e.g.* increasing the pasture size), and buying additional feed were commonly reported responses (Holman et al. 2021; Salmoral et al. 2020).

Changing crop types (*e.g.* from spring to winter crops), crop species or varieties can be successful adaptation strategies both in short and long term. Since the choice of crops and varieties are made before sowing/planting, such decisions can be considered as less of crisis management than *e.g.* increasing the irrigation or changing the harvest time and use of a specific crop. Regarding crop choice, grasses and grasslands may tolerate drought better than annual crops, but there seems to be a lack of information on how to combine species to optimise tolerance (Emadodin et al. 2021). Crops with deeper root systems are generally less sensitive to drought (van Oort et al. 2023). Hemp was found to be an interesting crop for adaptation to dryer conditions, since it can tolerate water shortage without over-exploiting soil water (Thevs and Nowotny 2023). Differences among crop varieties in drought tolerance have been found in *e.g.* spring (Nehe et al. 2023) and winter wheat (Statkevičiūtė et al. 2022), which is useful for the adaptation strategy to choose the most drought-tolerant varieties.

Several studies investigated modelling tools based *e.g.* on long-term weather and crop yield data to predict yield losses (Emadodin et al. 2021; Reinermann et al. 2019; van Oort et al. 2023; Webber et al. 2020). A combination of long-range seasonal weather forecasts and crop growth simulations confirmed that variations in weather and crop yield can be accurately foreseen by the used models, but showed that more precision is needed if such simulations are going to be useful for farmers’ decision-making regarding *e.g.* irrigation plans or choice of crops and varieties (Boas et al. 2023). Another study combined various meteorological sources, *i.e.* ground measurements, radar and satellite data in an effort to increase the predictive ability and precision of weather forecasts that can be used *e.g.* for management of irrigation systems in precision agriculture, which would be an adaptation at higher spatial and organizational levels than individual farms (Kepinska-Kasprzak and Struzik 2023).

Increasing the crop access to water by irrigation is reported as an efficient adaptation measure in drought situations and modelling the water balance to instruct irrigation management can improve water use efficiency (Campana et al. 2022; Shorachi et al. 2022; van Oort et al. 2023). Since the crop uptake of soil nitrogen is often reduced in dry conditions, due both to reduced crop growth and reduced nitrogen availability in dry soils, precision farming tools for crop nitrogen status can lead to fertilizer savings in dry years and improve nitrogen use efficiency by adapting fertilization to subsequent crops when higher levels of residual soil nitrogen are detected after a dry year (Klages et al. 2020).

Changed land use is another example of adaptation, and a study in south Germany found that the shade provided by solar panels in crop fields was beneficial for three out of four crops in 2018, indicating that solar panels can be an interesting tool for both climate change mitigation and adaptation to hotter and drier conditions (Trommsdorff et al. 2021).

### 3.2. Crisis management

One researcher-led crisis management influence diagram and one farmer-led crisis management influence diagram were constructed for the 2018 heat and drought event (Figure 4). The crisis management decisions identified were not necessarily motivated by expectations of a changing climate but were primarily motivated by an ongoing crisis.

Notably, farmers and scientists focused on different objectives. While cash crop and fodder yield were identified as basic and important farming objectives for both groups, low administrative burden was an important additional objective for farmers, whereas scientists focused on water and soil quality.

Moreover, the farmers did not highlight pests and diseases or beneficial organisms as uncertain factors as the scientists did.

It is also important to consider the role of agricultural advisors who may need to step in at short notice and give advice to farmers experiencing crises, such as the 2018 heat and drought event (see Box 1).

#### Box 1.

**The Role of Agricultural Advisors in Crisis Situations**

**Advisors as Key Intermediaries**

Agricultural advisors act as vital links between research, proven experience, and practical farming. Their role expands significantly during crises, requiring them to provide validated, context-sensitive solutions under regulatory and economic constraints.

**Institutional Diversity**

Advisory functions are embedded across various institutions—not only traditional advisory firms but also banks, insurance companies, public agencies, and commercial actors—each contributing to the broader support ecosystem.

**Case Study: The 2018 Drought in Sweden**

- **Early Signs**: Poor weather in 2017 led to late harvests and difficulties in crop establishment, already increasing advisory engagement.
- **Crisis Escalation**: In 2018, as drought conditions intensified, advisors provided guidance on harvest timing, irrigation, alternative fodder sources, and emergency strategies like forest grazing or importing feed.
- **Economic Stress**: Rising costs and reduced forage led to increased slaughter rates. Advisors, including financial institutions, played a key role in helping farms manage liquidity and credit access.
- **Investments**: Some farmers invested in irrigation or expanding usable land, enabled by financial advice and support.
- **Lasting Impacts**: The economic effects persisted for several years, sustaining demand for advisory services well beyond the drought itself.

**Psychosocial Support**

Crises also bring mental strain. Advisors are often among the first to identify and respond to psychosocial distress. Their presence can support farmers and families through stress, anxiety, and uncertainty—critical yet often under-recognized aspects of crisis resilience.

**Civic Preparedness and Social Networks**

Advisors, together with family, friends, and colleagues, form a social safety net. Strengthening civil society’s capacity to offer emotional and informational support is essential for effective long-term crisis management.

### 3.3 Climate change adaptation

Based on expert knowledge and insights from the scoping review, a researcher-led influence diagram on adaptation to climate change in crop production was constructed (Figure 5). Additionally, a corresponding influence diagram was constructed for farmers that was based on the discussions during the workshop.

Scientists and farmers focused on different objectives related to crop production (Figure 5). While both groups emphasised cash crops, fodder yield, and soil quality/fertility, scientists placed significant importance on water quality. In contrast, farmers identified low administrative burden as a critical objective that competes with climate change adaptation decision-making. Farmers did not prioritise pest and disease management as highly as scientists did. Additionally, scientists highlighted investment in irrigation and the monitoring of beneficial organisms and pests and diseases as important measures for climate change adaptation in crop production.

## 4. Discussion

This study presents an integrated analysis of the 2018 drought and heatwave in Sweden, drawing upon scientific literature, project scientistś insights, and local farmer knowledge to examine the impacts of extreme climatic events on crop production and climate change adaptation in agricultural systems. By comparing scientific and experiential perspectives, the study identified areas of alignment and disconnection between these knowledge systems, thereby highlighting critical gaps in adaptation knowledge. Employing a problem-feeding interdisciplinary and transdisciplinary approach, we considered not only the immediate effects of the drought and heat-wave but also broader challenges and opportunities posed by climate change.

### 4.1 Impacts and adaptation to the 2018 drought and heatwave in northern European agriculture

The 2018 drought and heatwave represented a severe climatic stressor for agriculture in northern Europe, including Sweden. Our review reveals that while numerous studies investigated the effects on specific crops or systems, the understanding of broader agricultural vulnerabilities remains fragmented, particularly regarding dose–response relationships and adaptation efficacy. Most studies reported yield reductions across a wide range of crops, including cereals, root vegetables, forage crops, and legumes. The few positive effects (e.g., dry soil facilitating certain field operations and reducing the need for post-harvest drying of grains, or fewer pests) were generally outweighed by significant negative impacts on crop productivity and farm economics (Holman et al. 2021).

#### 4.1.1 Evidence Gaps and Limitations in the scientific knowledge

A critical gap in the literature is the limited number of quantitative effect size studies that link specific levels of drought and heat exposure to yield outcomes. While long-term data sets have been used to confirm that the 2018 event was among the most damaging in decades, few studies quantified threshold responses or critical tipping points for different crops or systems. This constrains our ability to generalise findings or develop robust predictive models for future events.

Additionally, the lack of integrated pest and beneficial organism monitoring limits our understanding of how ecological processes are altered under compound climate stress. Despite isolated insights, such as the reduced impact of fungal disease in spring wheat in 2018 (Nehe et al. 2023) or the potential role of flower strips in supporting beneficial predators (Raderschall et al. 2022), systematic evaluation of pest dynamics during drought years remains scarce.

Another notable limitation is the geographic and sectoral distribution of the research. While studies span several northern European countries, there are few comparative analyses or regional syntheses that assess how local contexts (e.g., soil type, farming systems, irrigation availability) shape vulnerability and adaptive capacity. For Sweden specifically, detailed analyses of how different production regions responded to the crisis are lacking. Regarding plant traits and crop selection, some studies suggest that deeper-rooted crops and permanent pastures performed better under the extreme conditions of a drought-heatwave (Emadodin et al. 2021; van Oort et al. 2023). This also makes the job of advisors more difficult, since they rely on good quality scientific evidence to help support practitioners in times of crisis.

#### 4.1.2 Coping versus adaptation

Most reported responses to the 2018 event can be characterised as short-term coping strategies rather than forward-looking adaptation. These include shifting planting or harvest times, using straw for feed rather than soil incorporation, and reducing herd sizes. Such reactive measures, while essential during a crisis, may not contribute meaningfully to adaptation.

In contrast, proactive strategies, such as the use of alternative crop cultivars with improved drought and/or heat tolerance, investment in irrigation systems, and land use changes, were less frequently reported. There is still limited evidence on the large-scale feasibility of growing crops with high water use efficiency (*e.g.* hemp; Thevs and Nowotny 2023) or the utility of agrovoltaic systems (solar panels combined with crop cultivation; Trommsdorff et al. 2021). Furthermore, planning tools, such as seasonal forecasting and crop modelling, are in development but not yet widely adopted partly due to challenges in predictive accuracy, user integration and spatial resolution (Boas et al. 2023; Kepinska-Kasprzak and Struzik 2023). Innovative practices that offer dual benefits for climate mitigation and adaptation, such as agrovoltaics, deserve further exploration, especially in terms of their scalability and economic feasibility.

Most importantly, further research is needed to deepen our understanding of farmers’ motivations and drivers for engaging in climate change adaptation. This knowledge is crucial to identifying their communication needs and to developing evidence-based communication guidelines and policy instruments that are relevant, actionable, and needed. Such research should involve data collection from large and diverse samples of farmers to ensure representativeness and to enable segmentation based on emergent behavioural patterns and local context (soil type, access to irrigation, etc.), allowing for the tailoring of communication and policy strategies to the specific needs of different farmer groups (*cf*. Blennow et al. 2020; Palmér et al. 2024).

Finally, cross-national collaboration, particularly among countries in northern Europe, should be encouraged to facilitate comparative analyses across similar pedo-climatic zones and to enhance the transfer of effective adaptation strategies.

#### 4.1.3 Swedish context and opportunities

For Sweden, where both arable and livestock systems were heavily affected in 2018, there is a need to build on emerging knowledge from long-term datasets and cultivar trials. National agricultural policy could support on-farm experimentation and regional advisory systems that help tailor adaptation strategies to local conditions. Moreover, the 2018 event offers a rare but valuable empirical foundation for assessing both vulnerability and adaptive capacity, data that should be systematically gathered, archived, and analysed in preparation for future extremes.

### 4.2 Differing perspectives on crisis management and climate change adaptation

The influence diagrams developed in this study reveal key differences in how farmers and scientists perceive both crisis management and long-term adaptation to climate change in crop production (Figures 3 and 4). Notably, the crisis management diagrams, constructed separately for farmers and researchers, were deliberately limited to reflect decision-making in response to the acute 2018 drought and heatwave, rather than broader climate change considerations. This delineation was introduced at the request of the researchers to analytically separate short-term crisis responses from long-term adaptation planning. As a result, the absence of climate change as a direct motivator in these diagrams does not necessarily reflect a lack of awareness or concern among participants, but rather the defined scope of the exercise.

**Figure 3.**
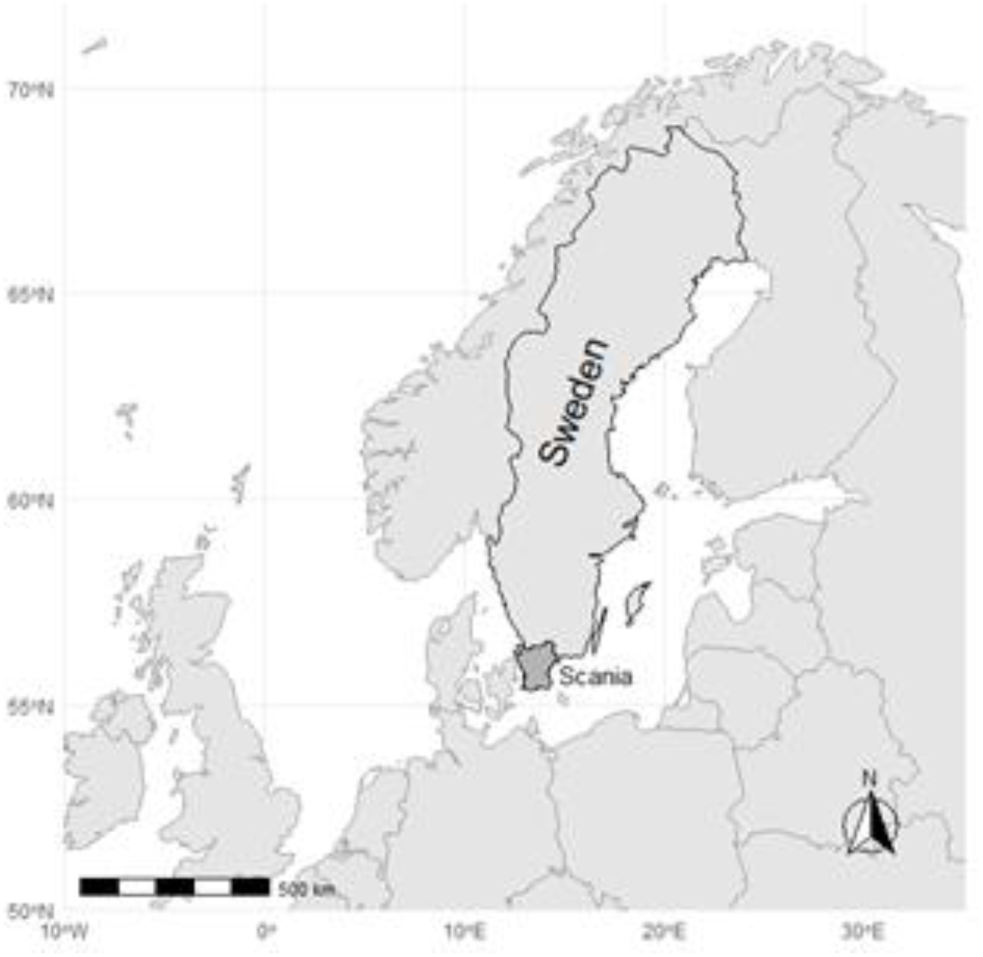
Map of northern Europe highlighting Sweden, with the location of Scania County. The map illustrates the geographical scope of the scoping review of scientific literature related to the 2018 heat and drought event, and marks Scania County where all participating farmers in the workshop are based. Created using Natural Earth data.

**Figure 4.**
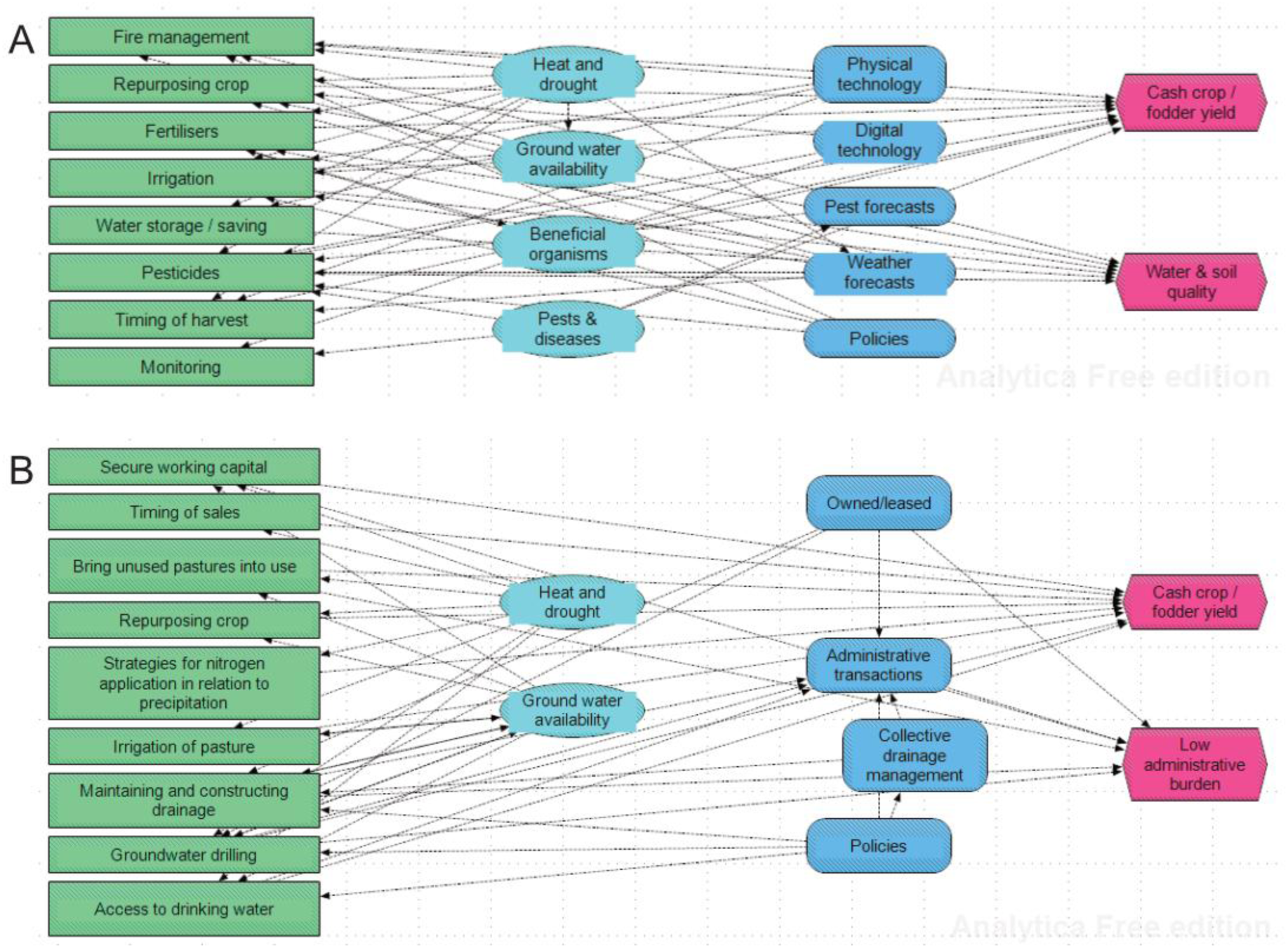
Influence diagrams depicting crisis management in crop production during the 2018 combined heat and drought event as seen by scientists (A) and farmers (B). The nodes represent decisions (green rectangles), chance variables that can be influenced by the decision-maker (light blue ovals), variables that cannot be directly influenced by the decision-maker (dark blue rounded rectangles), and objectives (red hexagons).

**Figure 5.**
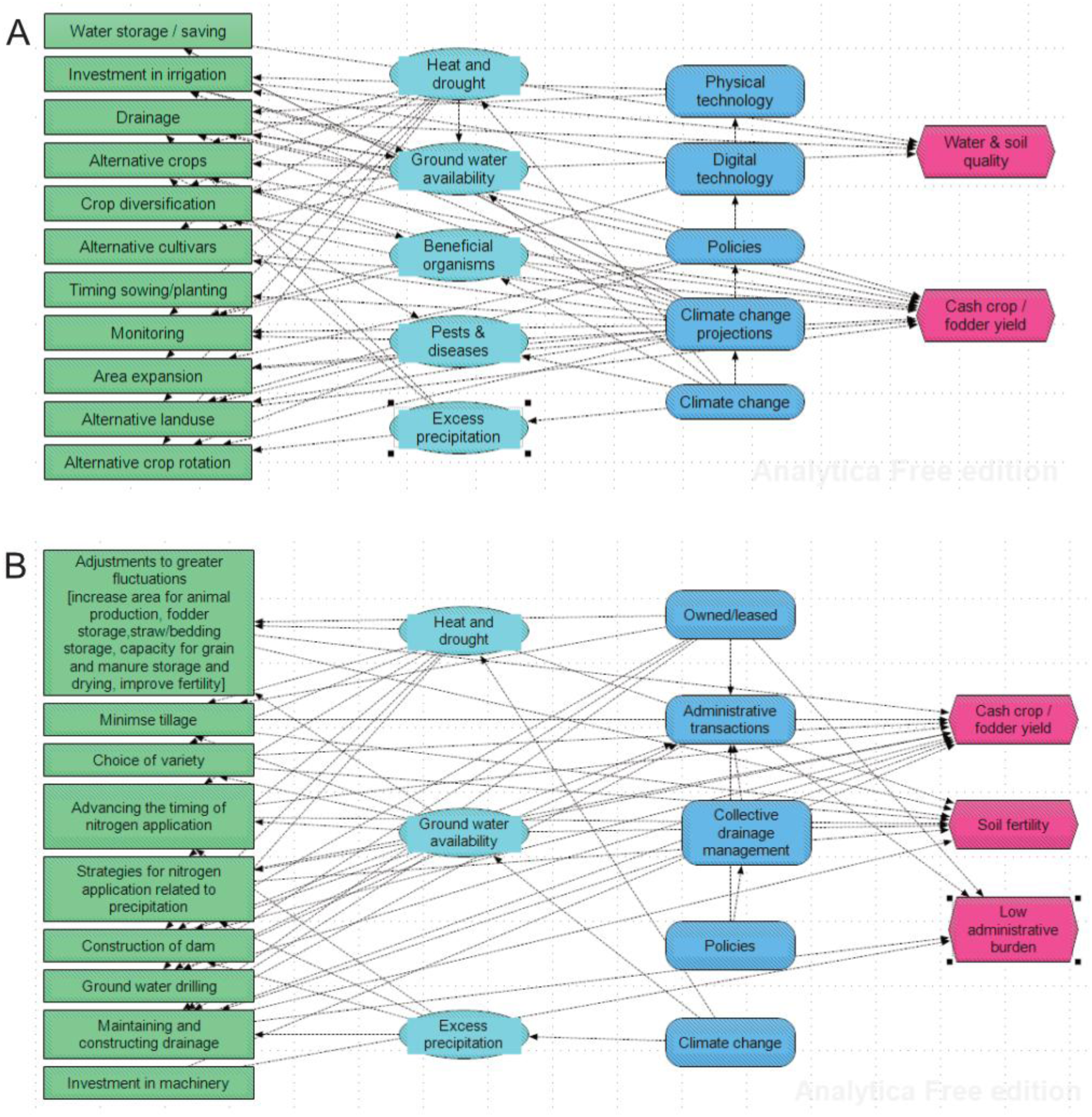
Influence diagrams depicting adaptation to climate change in crop production as seen by scientists (A) and farmers (B). The nodes represent decisions (green rectangles), chance variables that can be influenced by the decision-maker (light blue ovals), variables that cannot be directly influenced by the decision-maker (dark blue rounded rectangles), and objectives (red hexagons).

Even within this bounded focus on crisis management, differences emerged in the underlying objectives and priorities of the two groups (Figure 3). While both farmers and researchers emphasised protecting yields of cash crops and fodder, farmers additionally highlighted “low administrative burden” as a critical factor in decision-making. Researchers, by contrast, placed greater emphasis on soil and water quality, and identified biological uncertainties, such as pests and beneficial organisms, as important considerations, which were not prioritised by farmers. Considering the types of decisions listed in the influence diagrams, farmers’ decisions are sometimes more specific or elaborated than scientists, *e.g.* regarding fertilization and irrigation. This difference can be interpreted as a result of the tendency of farmers to reflect on details in their own specific situation while scientists are trained to identify factors that can be generalized across a wider scope. Furthermore, the farmers’ decisions included economic factors (securing capital, timing of sales) that were not part of the scientists’ influence diagram, and farmers emphasized socio-economic aspects (whether the land is owned or leased, administrative transactions and collective drainage management) rather than technologies and forecasts when discussing variables that cannot directly be influenced by the decision-maker. This tendency to link decisions on crisis management to socio-economic aspects might be related to the system-level view of farm operations held by farmers. This is reminiscent of findings by Carton et al. (2022) that post-harvest handling and possibilities to sell the products are part of farmers decisions on crop choice.

The influence diagrams on climate change adaptation further illustrate the divergence in priorities and constraints (Figure 4). Both groups recognised the importance of productive outcomes and soil fertility, but scientists proposed proactive adaptation measures such as investment in irrigation and monitoring of pests and beneficial organisms. Farmers, on the other hand, again stressed the burden of administrative processes as a significant barrier to implementing adaptive strategies. Their lower emphasis on pest and disease concerns may stem from region-specific experiences or from differing risk perceptions. Similar to the influence diagrams on crisis management, farmers decisions tended to be more detailed and specific than scientists (*e.g. “*maintaining and constructing drainage” *vs* “drainage”; “ground water drilling” *vs* “invest in irrigation”), and farmers included socioeconomic instead of technology-related variables that cannot be directly influenced by the decision-maker.

These findings, that highlight differences in what scientists and farmers emphasize in short-and long-term climate change adaptation, underscore the value of participatory, transdisciplinary methods in uncovering differences not only in proposed actions but in how goals and constraints are framed.

They highlight the need to co-develop adaptation strategies that integrate both scientific evidence and the lived realities of farming, while also acknowledging the complexity of decision-making in times of both crisis and transition. Collaborative efforts to identify and prioritize climate change adaptation strategies can combine farmers’ know-how and scientists know-why (Ingram et al. 2010). Moreover, farmers brought attention to variables beyond the farm, such as collective drainage management and administrative aspects of water management, probably driven by their system-level view on decision-making (Carton et al. 2022), which calls for multi-actor collaborations that include public authorities in order to facilitate the adaptation of agriculture to climate change. The influence diagrams serve as a powerful tool in this process, offering a visual representation of system-level thinking that can guide future communication, policy design, and adaptive planning.

### 4.3 Recommendations for future research and policy

Importantly, the results presented in this study also constitute an initial step toward designing a comprehensive survey targeting a broader population of farmers. By capturing the nuances of farmer experiences and decision-making, this exploratory work helps to formulate relevant and context-sensitive questions for future quantitative studies. The survey responses can be analysed to identify the drivers of adaptation decision-making, which in turn can inform the identification of farmer communication needs (*cf*. Blennow et al. 2020; Palmér et al. 2024). Such an approach strengthens the empirical basis for policymaking and communication strategies that are both evidence-based and aligned with on-the-ground realities.

Future research should also broaden the scope of effect size studies across a wide range of crops and agricultural systems to better characterise vulnerabilities and support the development of threshold-based risk assessments. Integrated systems research is particularly needed to examine crop performance in conjunction with ecological interactions, such as pest dynamics and pollinator activity, and farm-level economic outcomes under compound, interacting climate stressors.

In support of day-to-day and seasonal decision-making, tools that integrate long-range weather forecasts with crop growth models, irrigation scheduling, and precision fertilisation remain valuable.

However, these tools serve primarily to support short-term operational decisions and crisis response, rather than long-term adaptation. Therefore, it is equally critical to invest in research and tools explicitly aimed at supporting longer-term strategic adaptation, including planning for land use changes, crop diversification, and investment in resilient and preferably adaptable agricultural infrastructure. Research on long-term climate change adaptation should also embrace the multitude of societal perspectives on agricultural systems and not forget the importance of administrative burdens and the need for collective actions regarding *e.g.* water management.

Longitudinal studies are also necessary to monitor the uptake and effectiveness of adaptation measures over time. These should examine not only whether farmers are adopting recommended practices, but also how well these practices perform under varying climate conditions, what constraints or enablers affect their continued use, and how practitioners’ perceptions of risk, opportunity, and expected outcomes evolve in response to changing environmental and policy contexts.

## 5. Conclusions

This study advances understanding of how farmers in southern Sweden perceive and respond to both acute climate events and the broader challenge of climate change. Beyond highlighting divergences between scientific and local knowledge, it underscores the need for adaptation strategies that are context-aware and system-sensitive.

Rather than solely emphasizing short-term coping mechanisms, future research and policy must embrace the complexity of farming decisions, which are shaped by experiential knowledge, socio- economic constraints, and administrative systems. By visualizing and comparing farmer and researcher perspectives, here we demonstrate the value of participatory, transdisciplinary approaches in bridging knowledge gaps and fostering shared understanding.

Importantly, this work also establishes a basis for a large-scale, representative survey of Swedish farmers. Insights from that effort will be key to designing communication strategies that are not only evidence-based, but also actionable.

In a rapidly changing climate, supporting long-term adaptation requires more than technical solutions, it calls for inclusive processes that integrate diverse perspectives. Doing so will enhance adaptive capacity not only at the farm level, but also across the agricultural sector in northern Europe and beyond.

## Acknowledgements

We are especially thankful to the ten farmers who participated in the workshop, two of whom were also students in the Agricultural and Rural Management programme at SLU-Alnarp. Their insights and engagement were invaluable to the study. We also gratefully acknowledge the support of Åsa Ode and Jenny Casey Eriksson at the SLU-Alnarp library for their assistance with literature searches and reference management.

## Funding (information that explains whether and by whom the research was supported)

The research was funded by the Faculty of Landscape Architecture, Horticulture and Plant Production Sciences at the Swedish University of Agricultural Sciences (SLU) in Alnarp.

## Conflicts of interest/Competing interests (include appropriate disclosures)

The authors declare that they have no conflicts of interest.

## Ethics approval (include appropriate approvals or waivers)

This study involved the collection of personal data but did not include any sensitive personal data as defined under applicable regulations. In accordance with national and institutional guidelines, ethical approval was not required and was therefore not sought.

## Consent to participate

All participants took part in the workshop voluntarily and were informed about the purpose of the study. Informed consent was obtained from all individuals prior to their participation.

## Consent for publication

Not applicable. No identifying personal data are included in the publication.

## Availability of data and material (data transparency)

All data generated or analysed during this study are included in this published article.

## Code availability (software application or custom code)

Not applicable. No custom code or software was used in this study.

## Authors’ contributions

**Conceptualisation:** K.B.;

**Formal analysis**: K.B. – construction and development of influence diagrams;

**Investigation:** K.B., G.C., L.G.B., Å.L. – designing the review protocol and search strategy, and conducting the literature selection; G.C., L.G.B., Å.L. – conducting the analysis in the scoping review;

**Writing – original draft:** All authors;

**Writing – review & editing:** All authors;

**Visualization:** K.B. – development of the influence diagram;

**Project administration:** K.B. – overall project coordination; P.H. – organization of the farmer workshop;

**Funding acquisition**: All authors; K.B. led the grant application. All authors read and approved the final manuscript.

## References

A. Literature cited outside the review

Arreyndip NA (2021) Identifying agricultural disaster risk zones for future climate actions. PLoS ONE 16(12): e0260430. 10.1371/journal.pone.0260430

Bebber DP, Ramotowski MAT, Gurr SJ (2013) Crop pests and pathogens move polewards in a warming world. Nature Climate Change. 3:985–988. 10.1038/nclimate1990

Blennow K, Persson J (2021) To mitigate or adapt? Explaining why citizens responding to climate change favour the former. Land, 10(3):240. 10.3390/land10030240

Blennow K, Persson J, Tomé M, Hanewinkel M (2012) Climate change: believing and seeing implies adapting. PLOS ONE, 7(11):e50181. 10.1371/journal.pone.0050182

Blennow K, Persson J, Gonçalves L, et al. (2020) The role of beliefs, expectations and values in decision-making favoring climate change adaptation – implications for communications with European forest professionals Environmental Research Letters DOI:10.1088/1748-9326/abc2fa

Blennow K, Persson E, Persson J (2021) DeveLoP – A Rationale and Toolbox for Democratic Landscape Planning. Sustainability, 13:12955 10.3390/su132112055

Buras A, Rammig A, Zang CS (2020) Quantifying impacts of the 2018 drought on European ecosystems in comparison to 2003, Biogeosciences, 17, 1655–1672, 10.5194/bg-17-1655-2020

Carton N, Swiergiel W, Tidåker P, Röös E, Carlsson G (2022) On-farm experiments on cultivation of grain legumes for food – outcomes from a farmer–researcher collaboration. Renewable Agriculture and Food Systems, 37(5):457–467. doi:10.1017/S1742170522000102

Fischer EM, Sippel S, Knutti R (2021) Increasing probability of record-shattering climate extremes. Nature Climate Change, 11, 689–695. 10.1038/s41558-021-01092-9

Fischhoff B (1995) Risk perception and communication unplugged: Twenty years of progress. Risk Analysis, 15:137–145. 10.1111/j.1539-6924.1995.tb00308.x

Fischhoff B (2013) The sciences of science communication. Proceedings of the National Academy of Sciences 110: 14033–14039. 10.1073/pnas.1213273110

Gregory PJ, Johnson SN, Newton AC, Ingram JS (2009) Integrating pests and pathogens into the climate change/food security debate. Journal of Experimental Botany 60:2827–2838. DOI: 10.1093/jxb/erp080

Grusson Y, Wesström I, Joel A (2021) Impact of climate change on Swedish agriculture: Growing season rain deficit and irrigation need. Agricultural Water Management 251: 106858. 10.1016/j.agwat.2021.106858

Gvozdenac S, Dedic B, Mikic S, Ovuka J, Miladinovic D (2022) Impact of climate change on integrated pest management strategies. In: Climate Change and Agriculture, N. Benkeblia (Ed.). 10.1002/9781119789789.ch14

Haddaway NR, Macura B, Whaley P, and Pullin AS. 2017. ROSES for Systematic Map Protocols. Version 1.0. DOI: 10.6084/m9.figshare.5897284

Haddaway, N. R., Macura, B., Whaley, P. and Pullin, A. S. (2018). ROSES RepOrting standards for Systematic Evidence Syntheses: pro forma, flow-diagram and descriptive summary of the plan and conduct of environmental systematic reviews and systematic maps. Environmental Evidence, 7. 7. DOI:10.1186/s13750-018-0121-7

Hamilton-Webb A, Manning L, Naylor R, Conway (2017) The relationship between risk experience and risk response: a study of farmers and climate change. Journal of Risk Research, 20:11, 1379–1393 DOI: 10.1080/13669877.2016.1153506

Heeb L, Jenner E, Cock MJW (2019) Climate-smart pest management: building resilience of farms and landscapes to changing pest threats J. Pest Sci 92:951–969. 10.1007/s10340-019-01083-y

Ibrahim MA, Johansson M (2021) Attitudes to climate change adaptation in agriculture – A case study of Öland, Sweden. J. Rural Studies, 86:1:15. 10.1016/j.jrurstud.2021.05.024

Ingram J (2008) Are farmers in England equipped to meet the knowledge challenge of sustainable soil management? An analysis of farmer and advisor views. Journal of Environmental Management, 86:214–228. DOI: 10.1016/j.jenvman.2006.12.036

Ingram J (2014) Farmer-Scientist Knowledge Exchange. In: Thompson, P.B., Kaplan, D.M. (eds) Encyclopedia of Food and Agricultural Ethics. Springer, Dordrecht. 10.1007/978-94-007-0929-4_68

Ingram J, Fry P, Mathieu A (2010) Revealing different understandings of soil held by scientists and farmers in the context of soil protection and management. Land Use Policy, 27: 51–60. 10.1016/j.landusepol.2008.07.005.

JRC (2018) Joint Research Centre - MARS Bulletin - Crop monitoring in Europe, 26(7).

Labeyrie V, Renard D, Aumeeruddy-Thomas Y, et al. (2021) The role of crop diversity in climate change adaptation: insights from local observations to inform decision making in agriculture. Current Opinion in Environmental Sustainability, 51:15–23. 10.1016/j.cosust.2021.01.006

Lumina Decision Systems (2025). Analytica (Version 6.5.11.266) [Computer software]. https://www.lumina.com

Ouzzani M, Hammady H, Fedorowicz Z, Elmagarmid A (2016) Rayyan-a web and mobile app for systematic reviews. Syst Rev, 5(1):210. DOI: 10.1186/s13643-016-0384-4

Palmér C, Wallin A, Persson J, Aronsson M, Blennow K (2023) Effective communications on invasive alien species: Identifying communication needs of Swedish domestic garden owners. Journal of Environmental Management, 340:117995. 10.1016/j.jenvman.2023.117995

Persson J, Vareman N, Wallin A, Wahlberg L, Sahlin N-E (2019) Science and proven experience: a Swedish variety of evidence based medicine and a way to better risk analysis? Journal of Risk Research, 1–11 10.1080/13669877.2017.1409251

Persson J, Wallin A, Dewitt B, Wahlberg L (2022) Certainty and systematicity of practice-derived evidence matter for its relative importance in professional decision-making: Survey results on the role of proven experience in Swedish medicine, nursing, OT, dentistry, and dental hygiene. Journal of Nursing Studies Advances, 4:100074. 10.1016/j.ijnsa.2022.100074

Semenza J, Ploubidis GB, George LA (2011) Climate change and climate variability: personal motivation for adaptation and mitigation. Environmental Health 10:46. 10.1186/1476-069X-10-46

Sinclair VA, Mikkola J, Rantanen M, Räisänen J (2019) The summer 2018 heatwave in Finland, Weather, 74, 403–409, 10.1002/wea.3525.

Sjökvist E, Andersson M, Eklund A, Karlsson E, Norman M (2025) Klimatunderlag för klimat- och sårbarhetsanalyser. (In Swedish.) Klimatologi, Nr 74. Swedish Meteorological and Hydrological Institute. https://www.smhi.se/polopoly_fs/1.213224!/Klimatologi_74%20Klimatunderlag%20f%C3%B6r%20klimat-%20och%20s%C3%A5rbarhetsanalyser.pdf Accessed 21 February 2025.

Statistics Sweden (2019) Production of cereals, dried pulses, oilseed crops, potatoes and temporary grasses in 2018 Final statistics. Official Statistics of Sweden, JO 16 SM 1901. ISSN 1654-4137 https://www.scb.se/en/finding-statistics/statistics-by-subject-area/agriculture-forestry-and-fishery/agricultural-production/production-of-cereals-dried-pulses-and-oil-seeds/ Accessed 9 January 2025.

Swedish Board of Agriculture (2025) Storleksgrupp jordbruksmark, Variabel och År, Swedish Board of Agriculture, Jordbruksföretag och areal efter Län, År 2005-2022 (Agricultural Holdings and Area by County, Size Group of Agricultural Land, Variable and Year), Swedish Board Agric, Jönköping, Sweden. https://statistik.sjv.se. Accessed 9 January 2025.

Thorén H, Persson J (2013) The philosophy of interdisciplinarity: sustainability science and problem- feeding. Journal for General Philosophy of Science, 44:337–355. DOI:10.1007/s10838-013-9233-5

Toreti A, Belward A, Perez-Dominguez I, Naumann G, Luterbacher J, Cronie O, et al. (2019). The exceptional 2018 European water seesaw calls for action on adaptation. Earth’s Future, 7, 652–663. 10.1029/

de Toro A, Eckersten H, Libére N, von Rosen D (2015) Effects of extreme weather on yield of major arable crops in Sweden. SLU Report number: 086. Department of Energy and Tecnology, SLU. ISBN: 978-91-576-9323-5

Wilcke RAI, Kjellström E, Lin C, et al. (2020) The extremely warm summer of 2018 in Sweden – set in a historical context. Earth System Dynamics, 11:1107–1121. 10.5194/esd-11-1107-2020

Wiréhn L (2018) Nordic agriculture under climate change: A systematic review of challenges, opportunities and adaptation strategies for crop production, Land Use Policy, 77:63–74. 10.1016/j.landusepol.2018.04.059

Wood MD, Bostrom A, Bridges T, Linkov I (2012) Cognitive mapping tools: Review and risk management needs. Risk Analysis, 32:1333–1348. DOI: 10.1111/j.1539-6924.2011.01767.x

B. References of the systematic review

Ayling SM, Thompson J, Gray A, McEwen LJ (2021) Impact of reduced rainfall on above ground dry matter production of semi-natural grassland in south Gloucestershire, UK: A rainfall manipulation study. Front Environ Sci 9. 10.3389/fenvs.2021.686668

Boas T, Bogena HR, Ryu D et al. (2023) Seasonal soil moisture and crop yield prediction with fifth-generation seasonal forecasting system (SEAS5) long-range meteorological forecasts in a land surface modelling approach. Hydrol Earth Syst Sci 27:3143–3167. 10.5194/hess-27-3143-2023

Campana, PE, Lastanao P, Zainali S et al. (2022) Towards an operational irrigation management system for Sweden with a water–food–energy nexus perspective. Agric Water Manag 271:107734. 10.1016/j.agwat.2022.107734

Emadodin I, Corral DEF, Reinsch T et al. (2021) Climate change effects on temperate grassland and its implication for forage production: A case study from northern Germany. Agriculture 11:232. 10.3390/agriculture11030232

Holman IP, Hess TM, Rey D, Knox JW (2021) A multi-level framework for adaptation to drought within temperate agriculture. Front Environ Sci 8:589871. 10.3389/fenvs.2020.589871

Kepinska-Kasprzak M, Struzik P (2023) Monitoring of plant cultivation conditions using ground measurements and satellite products. Water 15:449. 10.3390/w15030449

Klages S, Heidecke C, Osterburg B (2020) The impact of agricultural production and policy on water quality during the dry year 2018, a case study from Germany. Water 12:1519. 10.3390/w12061519

Nehe A, Martinsson UD, Johansson E, Chawade A (2023) Genotype and environment interaction study shows fungal diseases and heat stress are detrimental to spring wheat production in Sweden. PLoS One 2023 18:e0285565. 10.1371/journal.pone.0285565

Raderschall C, Lundin O, Lindström SAM, Bommarco R (2022) Annual flower strips and honeybee hive supplementation differently affect arthropod guilds and ecosystem services in a mass-flowering crop. Agric Ecosyst Environ 326:107754. 10.1016/j.agee.2021.107754.

Reinermann S, Gessner U, Asam S et al. (2019). The effect of droughts on vegetation condition in Germany: An analysis based on two decades of satellite earth observation time series and crop yield statistics. Remote Sens 11:1783. 10.3390/rs11151783

Salmoral G, Ababio B, Holman IP (2020). Drought impacts, coping responses and adaptation in the UK Outdoor livestock sector: Insights to increase drought resilience. Land 9:202. 10.3390/land9060202

Shorachi M, Kumar V, Steele-Dunne SC (2022) Sentinel-1 SAR backscatter response to agricultural drought in the Netherlands. Remote Sens 14:2435. 10.3390/rs14102435

Statkevičiūtė G, Liatukas Ž, Cesevičienė J, et al. (2022) Impact of combined drought and heat stress and nitrogen on winter wheat productivity and end-use quality. Agronomy 12:1452. 10.3390/agronomy12061452

Thevs N, Nowotny R (2023) Water consumption of industrial hemp (*Cannabis sativa* L.) during dry growing seasons (2018–2022) in NE Germany. J Kulturpflanzen 75:173–184. 10.5073/JfK.2023.07-08.01

Trommsdorff M, Kang J, Reise C et al. (2021) Combining food and energy production: Design of an agrivoltaic system applied in arable and vegetable farming in Germany. Renew Sustain Energy Rev 140:110694. 10.1016/j.rser.2020.110694

van Oort PAJ, Timmermans NGH, Schils RLM, van Eekeren N (2023) Recent weather extremes and their impact on crop yields of the Netherlands. Eur J Agron 142:126662. 10.1016/j.eja.2022.126662

Webber H, Lischeid G, Sommer M et al. (2020) No perfect storm for crop yield failure in Germany. Environ Res Lett 15:104012. 10.1088/1748-9326/aba2a4

